# MEDIPIPE: an automated and comprehensive pipeline for cfMeDIP-seq data quality control and analysis

**DOI:** 10.1101/2023.02.28.530481

**Authors:** Yong Zeng, Ye Wenbin, Eric Y. Stutheit-Zhao, Ming Han, Scott V. Bratman, Trevor J. Pugh, Housheng Hansen He

## Abstract

**Summary:** cell-free methylated DNA immunoprecipitation and high-throughput sequencing (cfMeDIP-seq) has emerged as a promising non-invasive technology to detect cancers and monitor treatments. Several bioinformatics tools are available for cfMeDIP-seq data analysis. However, an easy to implement and flexible pipeline, particularly, for large-scale cfMeDIP-seq profiling, is still lacking. Here we present the MEDIPIPE, which provides a one-stop solution for cfMeDIP-seq data quality control, methylation quantification and sample aggregation. The major advantages of MEDIPIPE are: 1) it is easy to implement and reproduce with automatically deployed execution environments; 2) it can handle different experimental settings with a single input configuration file; 3) it is computationally efficient for large-scale cfMeDIP-seq profiling data analysis and aggregation.

**Availability and implementation:** This pipeline is an open-source software under the MIT license and it is freely available at https://github.com/yzeng-lol/MEDIPIPE.

**Supplementary information:** Supplementary data are appended.

## 1 Introduction

cell-free DNA (cfDNA) in blood has been a promising analyte for cancer prognosis and treatment monitoring (Corcoran and Chabner, 2018). Next-generation sequencing (NGS)-based technologies have also been tailored to identify cfDNA genomic and epigenomic signatures associated with cancer phenotypes. Among them, the epigenomic profiling through cell-free methylated DNA immunoprecipitation and high-throughput sequencing (cfMeDIP-seq) (Shen *et al*., 2018) has proven capable of ultrasensitive tumor detection and classification, particularly in the scenario of early-stage cancer or minimal tumor residual after treatments (Shen *et al*., 2018; Burgener *et al*., 2021; Nassiri *et al*., 2020). Refined cfMeDIP-seq protocols incorporating spike-in controls and/or unique molecular identifiers (UMIs) also offer improved capabilities for batch effect correction and error-suppression (Shen *et al*., 2019; Burgener *et al*., 2021). We have also been witnessing an increased number of large-scale cfMeDIP-seq profiling studies conducted in different cancer types (Shen *et al*., 2018; Liu *et al*., 2021; Burgener *et al*., 2021; Nuzzo *et al*., 2020; Nassiri *et al*., 2020; Chen *et al*., 2022). However, an easy-to-implement and flexible pipeline for large-scale cfMeDIP-seq profiling data processing and analysis is still lacking in the field.

Here we present an open-source pipeline, MEDIPIPE, to provide an automated and comprehensive solution for cfMeDIP-seq data quality control (QC), methylation quantification and sample aggregation. This pipeline was developed using Snakemake (Mölder *et al*., 2021), ensuring all dependencies are automatically installed and execution is seamless and reproducible. This pipeline is able to handle various experimental settings, such as sequence layout and whether spike-in controls and/or UMI were added, via specifying a single input configuration file. Moreover, it can efficiently deal with large-scale cfMeDIP-seq profiling data on high performance computing clusters, because all independent steps for individual samples are run in parallel.

## 2 Pipeline description

MEDIPIPE consists of four modules, which starts with parsing the customized input configuration file, and subsequently conducting corresponding workflows (**Fig. 1 and Supplementary Fig. S1**). In general, the raw cfMeDIP-seq sequencing reads will be preprocessed, aligned, quantified and quality assessed per sample in the first three modules. Then, users can activate the final module to generate aggregated QC reports and quantification matrices.

**Fig.1.**
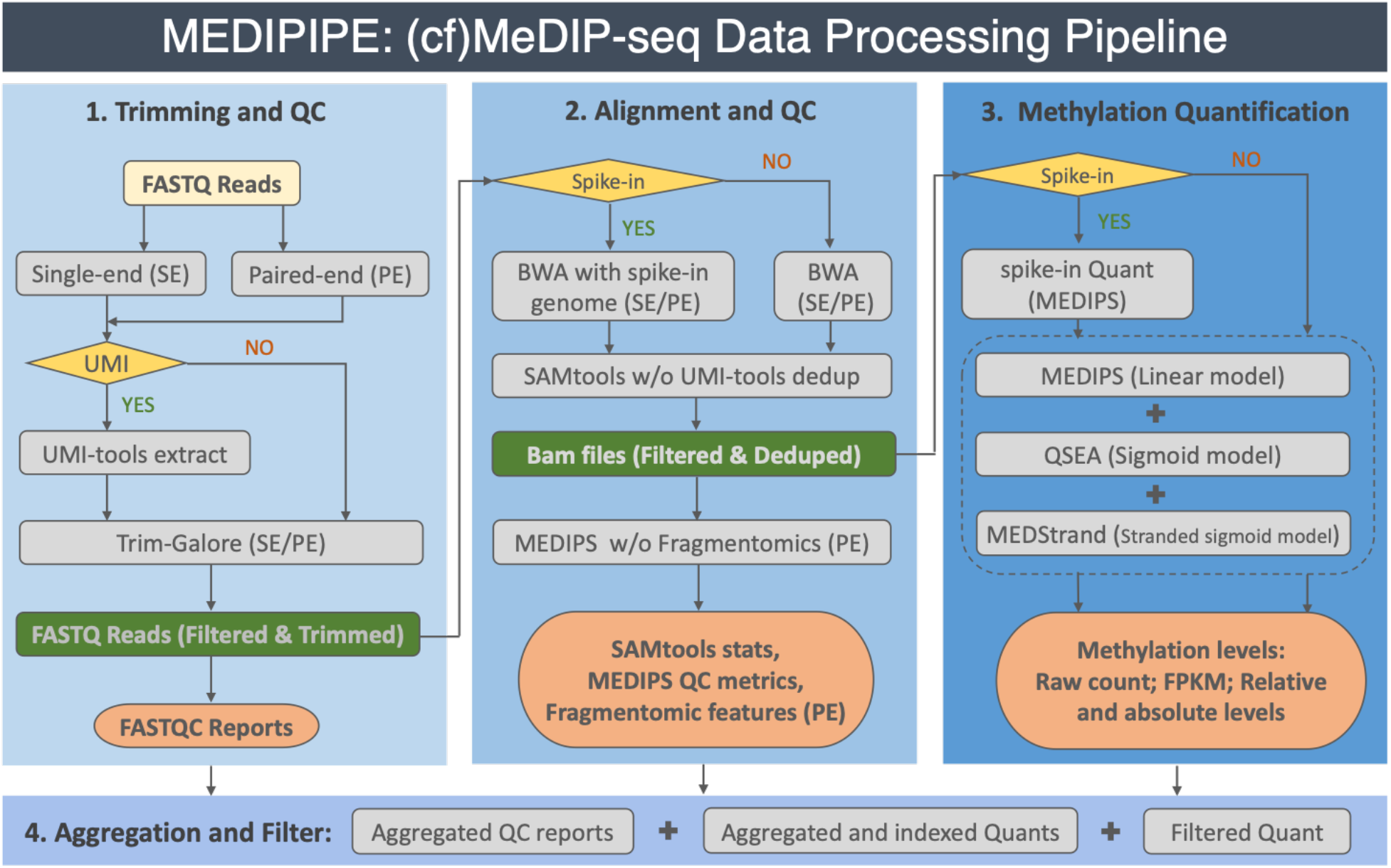
Flowchart of the MEDIPIPE pipeline. UMI, unique molecular identifiers; QC, quality control.

### 2.1 Input of pipeline

A single input configuration file in YAML format is required for successfully running this pipeline, which specifies the paths to the working environment, genome references table, samples’ sequencing data, and aggregation tables, as well as parameters for different experimental settings. Instructions and a detailed template are included in the repository. Notably, the sequencing reads can be either single-end or paired-end gzip-compressed FASTQ files, and multiple FASTQ files (e.g. multiple sequencing runs) for the same biological sample can be merged by the pipeline as well. We also provide shell scripts for automatically downloading ENCODE pre-build genome references (e.g. BWA index and annotated regions) (Luo *et al*., 2020), or building customized genome references (e.g. when spike-in sequences were added to the primary genome).

### 2.2 Reads trimming and QC

Trim Galore (https://www.bioinformatics.babraham.ac.uk/projects/trim_galore/) is employed to trim adapter sequences and low-quality bases of raw reads. If UMI barcodes are added into the library to suppress deduplication error, UMI-tools (Smith *et al*., 2017) will be executed prior to Trim Galore to extract barcodes and remove those reads which failed to match the barcode pattern. Lastly, FastQC (https://www.bioinformatics.babraham.ac.uk/projects/fastqc/) is applied to check the qualities of both raw and preprocessed reads. All of these steps can be executed in either singleend or paired-end mode (**Fig. 1**).

### 2.3 Reads alignment and QC

Next, the preprocessed reads will be mapped back to corresponding genome(s) via BWA-MEM (Li, 2013). For the scenario of reads coming along with spike-in controls, we recommend appending spike-in sequences to the primary genome for alignment simultaneously. Aligned spikein reads can be extracted for separate quantity assessment and quantification, which is specified in the configuration file. Then, SAMtools (Li *et al*., 2009) is applied to filter out unmapped reads and secondary alignments, as well as improperly paired mates for paired-end reads. Duplicated reads will be removed either by SAMtools markdup (Li *et al*., 2009) or UMI-tools dedup (Smith *et al*., 2017), depending on whether UMI barcodes were added or not (**Fig. 1**). Specific to paired-end reads, the fragment size will be estimated by CollectInsertSizeMetrics from Picard tool kit (http://broadinstitute.github.io/picard/), and the fragmentation profiles, which is defined as the ratio of short to long fragments in consecutive windows (Cristiano *et al*., 2019), will be computed at 1 megabase pair (Mb) and 5Mb resolution as well. Lastly, QC metrics derived from the aligned reads, including BAM files statistics via SAMtools (Li *et al*., 2009) and cfMeDIP-seq saturation, coverage, and enrichment scores via MEDIPS (Lienhard *et al*., 2014), are collected.

### 2.4 Methylation quantification

The cfMeDIP-seq offers higher coverage of methylated CpG dinucleotides throughout the genome with much lower cost compared to bisulfite-based methods, however, the absolute methylation levels can only be estimated using a computational model for this enrichment based profiling method. MEDIPIPE applies three successively developed methods for relative and absolute methylation estimation (**Supplementary Table S1**): MEDIPS (Lienhard *et al*., 2014), QSEA (blind) (Lienhard *et al*., 2017) and MeDStrand (Xu *et al*., 2018). These methods were designed to eliminate CpG density bias with a linear regression model, sigmoidal model with empirical knowledge, and stranded sigmoid model, respectively. This allows the users to choose the quantifications as they need and enables a comprehensive comparison among different estimations. If there are spike-in controls, MEDIPS (Lienhard *et al*., 2014) is used to separately quantify the methylation levels for them as well.

### 2.5 Aggregation and filtering

Since many cfMeDIP-seq profiling projects generate data for a large number of samples in multiple groups or batches, the final module of MEDIPIPE can aggregate QC metrics and methylation matrices across all samples for each such project. Namely, first, all QC reports generated by FASTQC, SAMtools, Picard and MEDIPS are aggregated by either MultiQC (Ewels *et al*., 2016) or an embedded script. On top of that, a summarized QC report in HTML format with selected QC metrics is generated, allowing users to interactively examine sample QC metrics across different groups or batches (**Supplementary Fig. S2**). Meanwhile, different methylation quantifications are aggregated into corresponding TAB-delimited bin-sample TXT files, as well as uniformly filtered out sex chromosomes, mitochondrial chromosome, and ENCODE blacklist regions (Amemiya *et al*., 2019). Both original and filtered aggregated quantification files are indexed by Tabix (Li, 2011), enabling rapid retrieval of data for genomic regions of interest for downstream analysis.

### 2.6 Outputs of pipeline

Since many All outputs of MEDIPIPE are organized in corresponding folders. Specifically, the main output files per sample can be grouped into four categories: QC reports, aligned BAM files, fragmentomic features (for paired-end reads only) and methylation quantifications (**Supplementary Table. S1**). Moreover, the aggregated outputs include multiQC reports, aggregated QC reports and the aggregated methylation quantification before and after filtering (**Supplementary Table. S2**)

## 3 Implementation

MEDIPIPE was developed with Snakemake following a clean, modular and robust design in accordance with best practice coding standards; other specialized tools can be easily added in the future. Detailed instructions on how to install and run this pipeline are presented in (https://github.com/yzeng-lol/MEDIPIPE). This pipeline is highly flexible thanks to the input configuration file, which comes along with options and parameters for different experimental settings and analyses. The pipeline can be run locally or submitted to the high performance computing (HPC) for efficient scheduling within a multiprocessor environment. We had also successfully run it with a published cfMeDIP-seq dataset (Nassiri *et al*., 2020), which consists of 163 samples from 6 brain cancer subtypes, to get aggregated quantifications and QC reports (**Supplementary Fig. S2).** Lastly, although this pipeline was originally developed for cfMeDIP-seq data, it can also be used for the analysis of methylated DNA immunoprecipitation followed by sequencing (MeDIP-seq) data.

## Acknowledgement

We like to thank Sasha Main, Emma Bell, Nicholas Cheng, Althaf Singhawansa and Justin Burgener for acting as the pipeline’s beta-tester and their feedback. This is a pilot project for the initiative of cell-free Multiomics Data Coordination Centre (cfMOS-DCC), we would also like to thank the cfMOS-DCC members for sharing their cfMeDIP-seq data for pipeline test and finetuning.

## Supplementary Data

### Supplementary Figures

**Supplementary Fig. S1.**
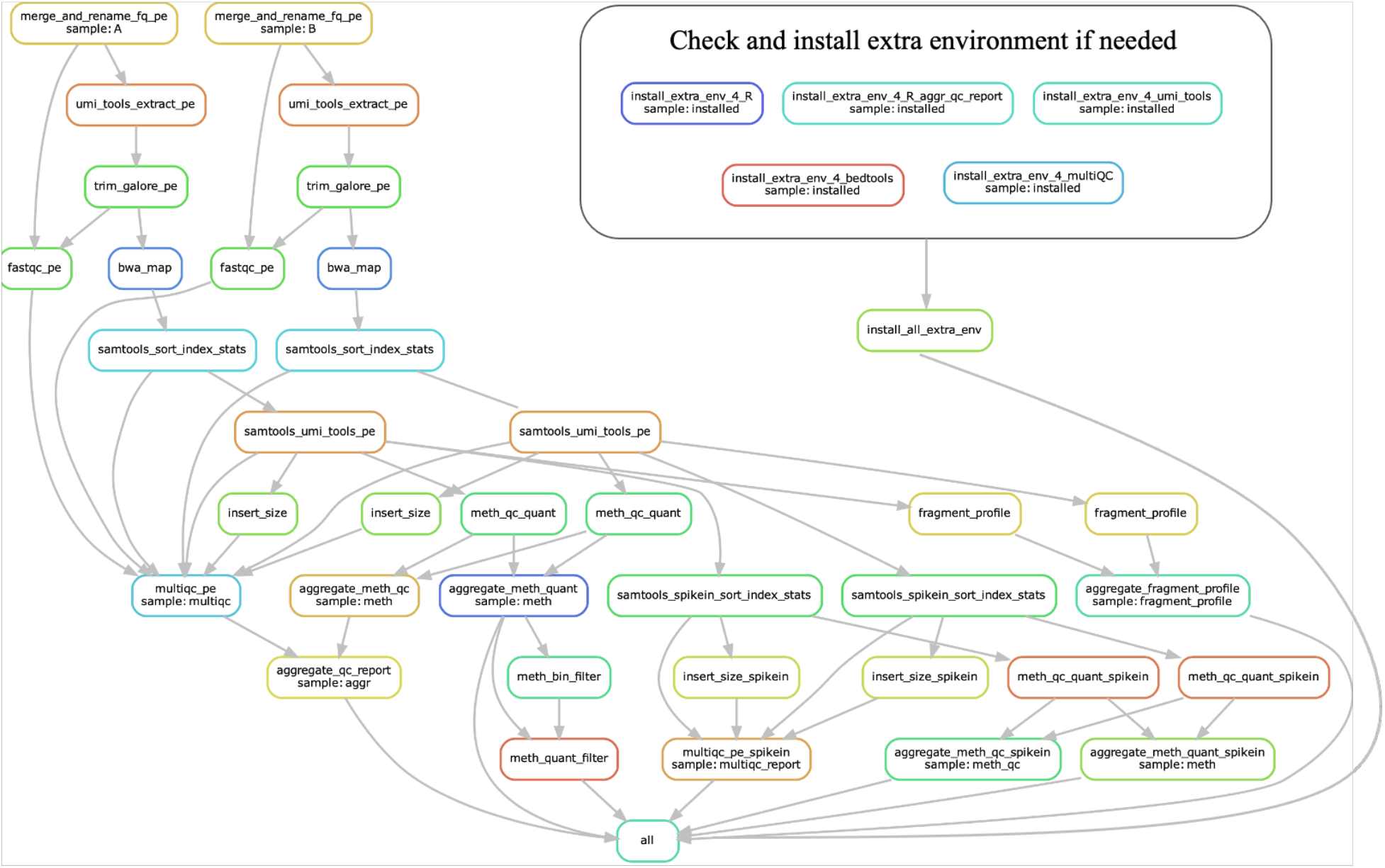
Example of directed acyclic graph (DAG) for MEDIPIPE dealing with two paired-end cfMeDIP-seq samples with UMI barcodes and spike-in controls.

**Supplementary Fig. S2.**
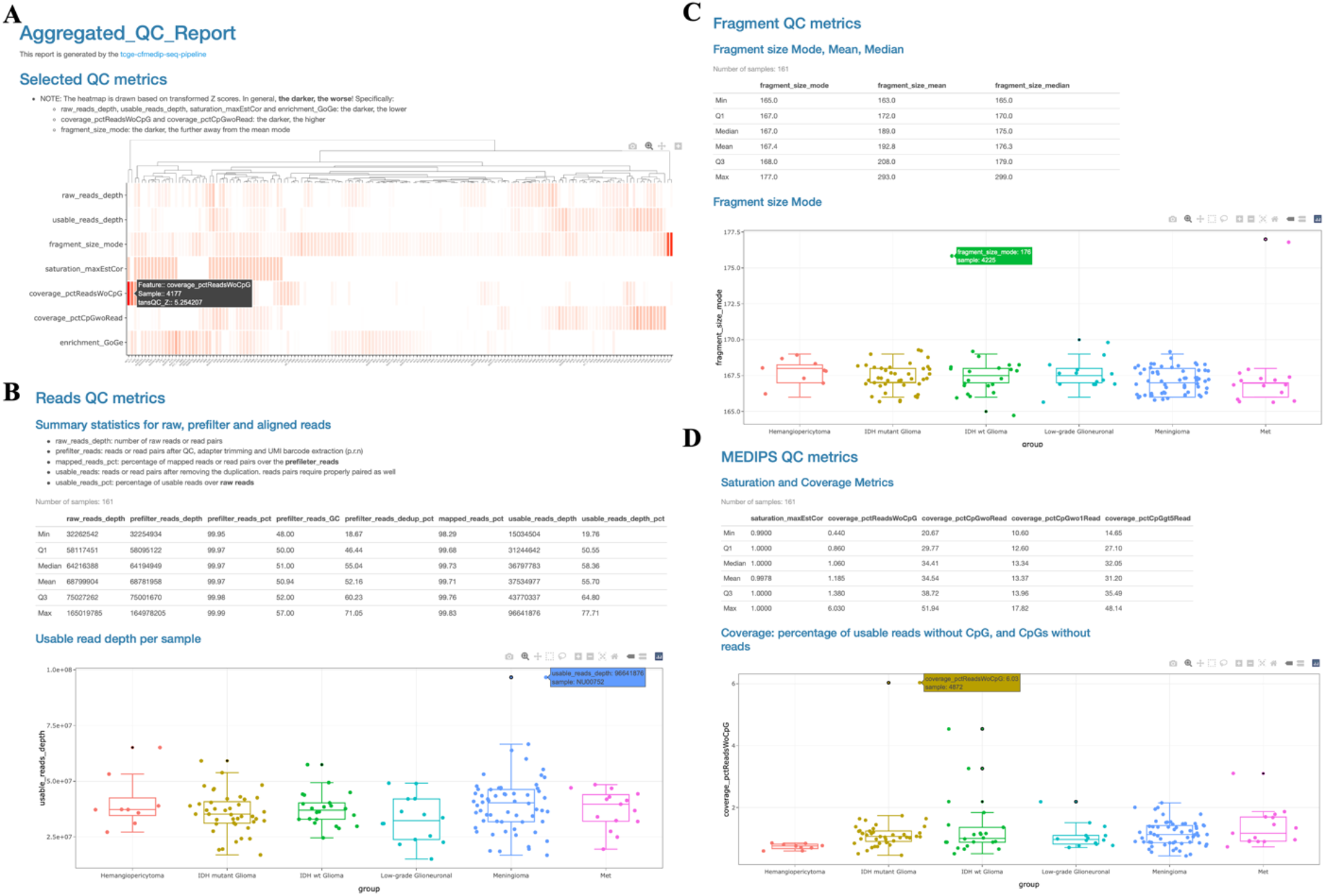
Example of an aggregated QC report, which is interactive in HTML format. **(A)** Summary heatmap based on transformed Z scores of selected QC metrics. **(B, C, D)** Summary statistics for Reads, Fragment and MEDIPS QC metrics, as well as boxplots with jittered samples per groups for corresponding QC metrics.

### Supplementary Tables

**Supplementary Table. S1:**
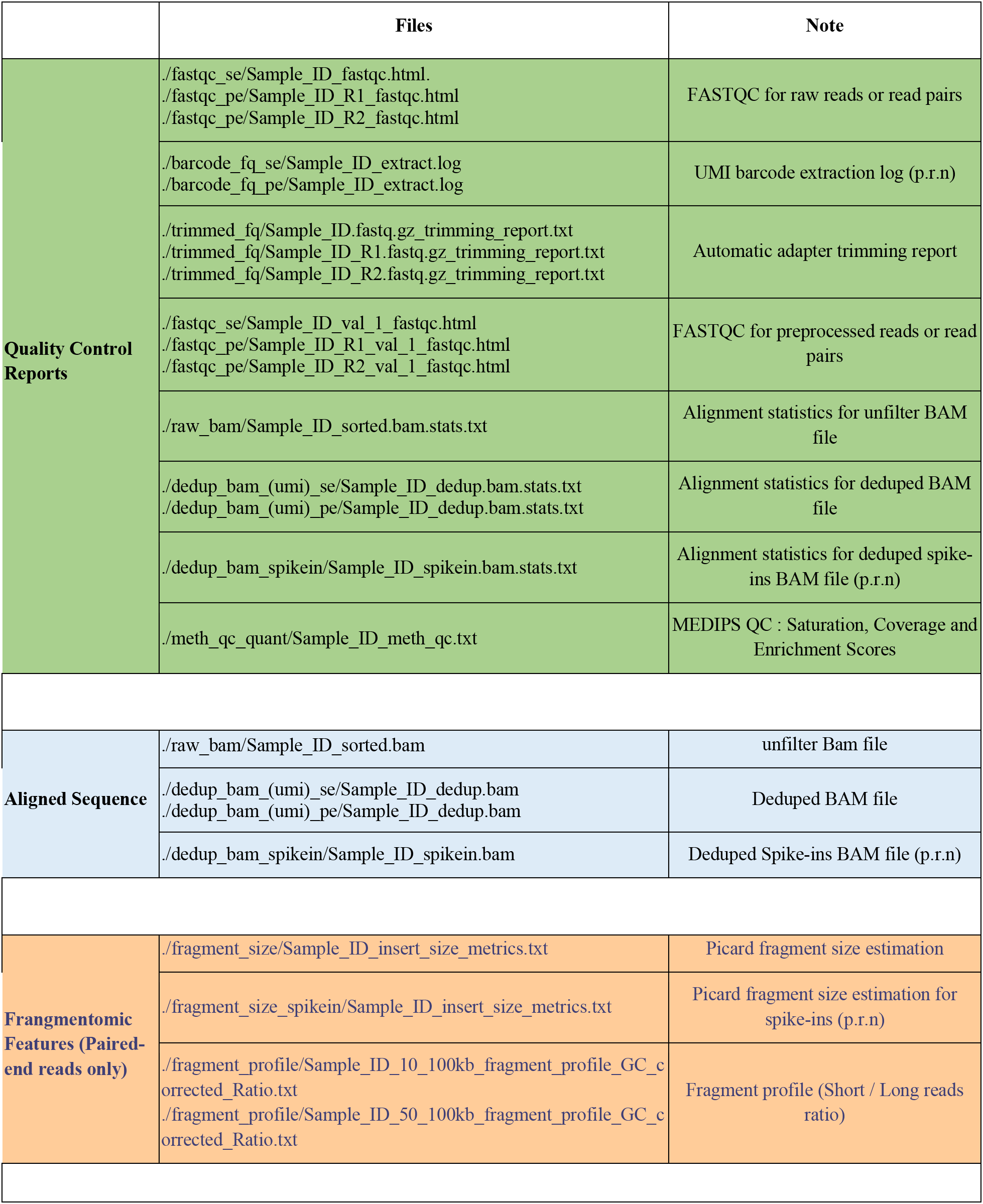
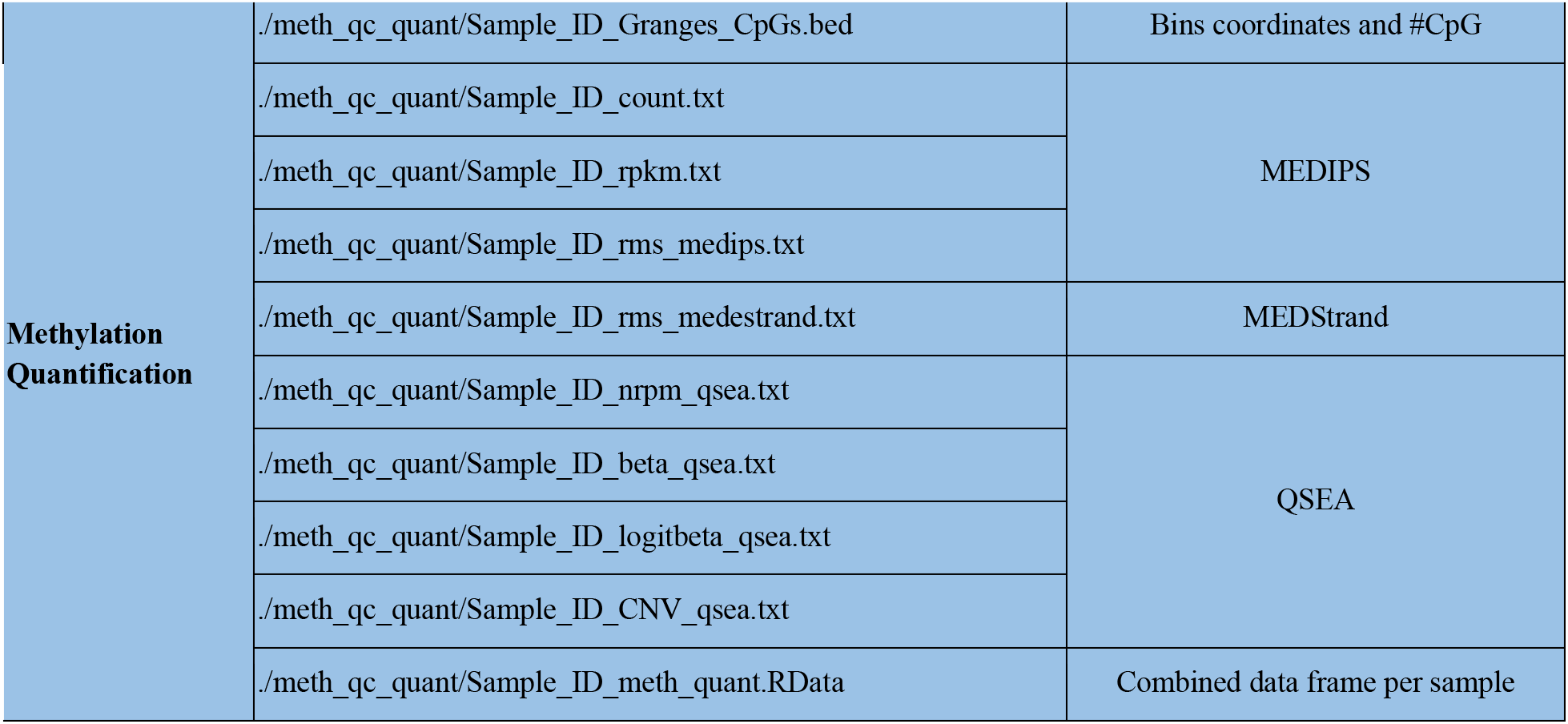
Output files per individual samples.

**Supplementary Table. S2:** Output files for aggregated samples

|  | Files | Note |
| --- | --- | --- |
| **Aggregated Quality Control Reports** | ./aggregated/QC_se/multiqc_data<br>./aggregated/QC_pe/multiqc_data | MultiQC data |
|  | ./aggregated/QC_se/multiqc_report.html<br>./aggregated/QC_pe/multiqc_report.html | MultiQC HTML report |
|  | ./aggregated/meth_qc.txt | Combined MEDIPS QC metrics |
|  | ./aggregated/aggr_qc_report.csv | Aggregated selected QC metrics |
|  | ./aggregated/aggr_qc_report.html | Aggregated QC HTML report |
| **Aggregated methylation quantification (Indexed with Tabix)** | ./aggregated/meth_bin.bed | Bins coordinates and #CpG |
|  | ./aggregated/meth_count.txt.gz | MEDIPS |
|  | ./aggregated/meth_rpkms.txt.gz |  |
|  | ./aggregated/meth_rms_medips.txt.gz |  |
|  | ./aggregated/meth_rms_medstrand.txt.gz | MEDStrand |
|  | ./aggregated/meth_nrpm_qsea.txt.gz | QSEA |
|  | ./aggregated/meth_beta_qsea.txt.gz |  |
|  | ./aggregated/meth_logitbeta_qsea.txt.gz |  |
|  | ./aggregated/meth_CNV_qsea.txt.gz |  |
|  | ./aggregated/meth_meth_quant.RData | Combined data frame per sample |
| **Aggregated methylation quantification and filtered out sex chromosomes, chromosome mitochondria and ENCODE blacklist regions (Indexed with Tabix)** | ./autos_bfilt/meth_autos_bfilt_bin.bed | Bins coordinates and #CpG |
|  | ./autos_bfilt/meth_count_autos_bfilt.txt.gz | MEDIPS |
|  | ./autos_bfilt/meth_rpkms_autos_bfilt.txt.gz |  |
|  | ./autos_bfilt/meth_rms_medips_autos_bfilt.txt.gz |  |
|  | ./autos_bfilt/meth_rms_medstrand_autos_bfilt.txt.gz | MEDStrand |
|  | ./autos_bfilt/meth_nrpm_qsea_autos_bfilt.txt.gz | QSEA |
|  | ./autos_bfilt/meth_beta_qsea_autos_bfilt.txt.gz |  |
|  | ./autos_bfilt/meth_logitbeta_qsea_autos_bfilt.txt.gz |  |
|  | ./autos_bfilt/meth_CNV_qsea_autos_bfilt.txt.gz |  |

